# The endogenous modulation of visual plasticity in human adults

**DOI:** 10.1101/2022.10.27.513994

**Authors:** Yiya Chen, Yige Gao, Zhifen He, Zhouyuan Sun, Yu Mao, Robert F. Hess, Peng Zhang, Jiawei Zhou

## Abstract

The adult human visual system can exhibit a degree of neuroplastic change under the right conditions which has implications for future treatments to recover vision loss that have occurred in infancy. The exogenous factors that promote neuroplastic change have been the focus of human psychophysical, electrophysiological and neuroimaging research over the last decade. What has not been considered is the importance of endogenous factors. In this study we modulate the neural oscillations that determines the internal neural state to demonstrate that endogenous factors play a critical role in not only baseline contrast sensitivity but also the extent to which the adult visual system can undergo neuroplastic change in binocular balance.

## Introduction

Recent studies have shown that when one eye is deprived of its input for a short period of time (30 min to 2.5 hrs), visual brain mechanisms undergo a neuroplastic change that results in not only a change in visual sensitivity but also interocular balance (i.e. ocular dominance) (Lunghi, Burr et al., 2011, Zhou, Clavagnier et al., 2013). A number of results suggest that this neuroplastic change mainly involves V1 (the primary visual cortex), these include intrinsic optical imaging in monkeys (Begum and Tso, 2015), and magnetoencephalography (MEG) (Chadnova, Reynaud et al., 2017), electroencephalogram (EEG) studies (Lunghi, Berchicci et al., 2015, Zhou, Baker et al., 2015), functional magnetic resonance imaging (fMRI) studies (Binda, Kurzawski et al., 2018), magnetic resonance spectroscopy (MRS) studies(Lunghi, Emir et al., 2015) and psychophysics studies (Zhou, Reynaud et al., 2014) in human adults. The role played by a number of exogenous factors have been investigated, namely image properties (Zhou, Reynaud, 2014, Zhou, Reynaud et al., 2017), exercise (Lunghi and Sale, 2015), visual pathology (Lunghi, Galli-Resta et al., 2019), body mass index (Lunghi, Daniele et al., 2019). What is not known is whether there is a role played by endogenous factors in modulating this neuroplastic change.

It is well known that there are characteristic differences in resting-state brain activity in the absence of visual stimulation, for example, when the two eyes are open versus closed in the dark, there is a significant decrease in occipital alpha oscillations called the Berger effect. There are power and coherence changes of a broad spectrum in the Δ, θ, α_1_, α_2_, β_1_, β_2_ and γ frequency bands that are presumed to be correlates of the switching of involuntary preliminary attention from internally directed attention specific for the eyes closed state to externally directed attention specific for the eyes open state (Boytsova and Danko, 2010). FMRI studies have shown that eyes open rest conditions are associated with larger activation of the visual cortex but smaller activation of the lateral geniculate nucleus (Marx, Deutschlander et al., 2004). In the eyes closed state, activations of the ocular motor related brain areas are larger, including the prefrontal eye fields, parietal and frontal eye fields, cerebellar vermis, thalamus and basal ganglia (Marx, Deutschlander, 2004). It has been argued that eye closure can alter the processing mode of the sensory system by decoupling geniculostriate processing in favor of enhanced thalamocortical coupling in non-visual brain areas (Brodoehl, Klingner et al., 2015). While it remains an important question whether this type of intrinsic regulation can modulate visual sensitivity and visual plasticity, it is a difficult question to answer because any comparison between eyes open and eyes closed conditions necessarily excludes the ability to measure visual sensitivity/plasticity using external visual inputs. Using the ocular dominance plasticity paradigm described above where one eye is deprived of its visual input for a short period of time, we have been able to assess the role of internal state (comparing eyes open with eyes close behind the deprivation patch) for visual sensitivity and visual plasticity in adult humans. In particular, we directly compare the monocular deprivation effects, as assessed by changes in EEG power, steady-state visually evoked potentials (SSVEP), and contrast sensitivity, during the period when the patched eye is kept either open or closed *behind* the patch. We derive contrast gain and ocular dominance changes that occur as a result of patching one eye for a 2.5-hour period when the eye *behind* the patch is either open or closed. The results show that this neuroplastic change can be modulated by the internal state in the absence of visual stimulation and is greater when the eye behind the occluded in kept open (i.e., eye open state).

## Results

### 1. The immediate effects of open v.s. close of the patched eye on intrinsic neural oscillations and the unpatched eye’s sensitivity

To investigate whether there is an internal state difference in intrinsic neural oscillations when only one eye is open or closed, we recruited 20 normal adults and measured the amplitudes of their alpha oscillations at the resting state with the two eyes closed, two eyes open, monocular patching with the patched eye (PE) open and monocular patching with the PE closed. Figure 1a shows the amplitude spectrum averaged across subjects for the 4 conditions. Figure 1b shows the average amplitude at the alpha peak frequency. Similar to previous reports (Boytsova and Danko, 2010), alpha amplitude was significantly lower when two eyes are open compared to closed (t(19) = - 2.272, *p* = 0.035, 2-tailed paired samples t-test with Holm-Bonferroni correction(Holm, 1979), number of comparisons k = 2). Importantly, in the monocular patching conditions, the alpha amplitude was significantly lower when the PE remaining open as compared to the PE being kept closed (t(19) = -3.944, *p* < 0.001, with Holm-Bonferroni correction, k = 2).

**Figure 1.**
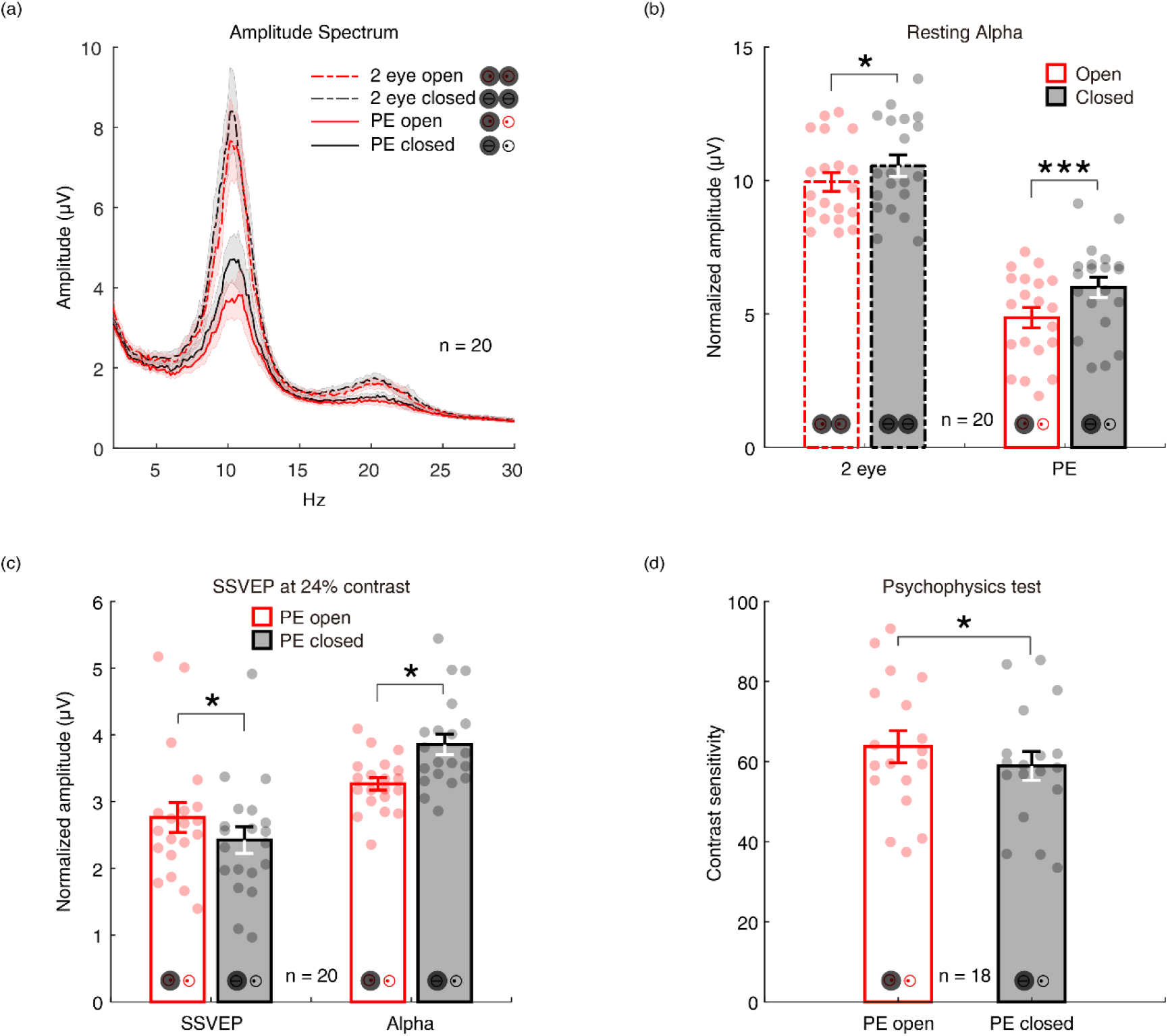
The immediate effects of open v.s. close of the patched eye. (a) the amplitude spectrum averaged across subjects between 2 to 30 Hz for 4 conditions (two eyes open: red dashed line; two eyes closed: black dashed lines; PE open: red solid line; PE closed: black solid line). Shaded area indicates one standard error of the mean (SEM) across 20 subjects; (b) the normalized amplitude at the alpha peak frequency. Each dot represents one subject. Error bars represent one SEM across 20 subjects. (c) The average amplitude of SSVEP and peak alpha. Each dot represents each observer. Error bars represent one SEM across 20 subjects; (d) the contrast sensitivity of UPE. Each dot represents one observer. Error bars represent one SEM across 18 subjects. *: *p* < 0.05; ***: *p* < 0.001.

We further measured SSVEP to a sinewave plaid at 24% contrast counterphase flickering at 7.5 Hz, and the contrast sensitivity of the unpatched eye (UPE) for both PE open and closed conditions. The results in Figure 1c show stronger SSVEP (t(19) = 2.737, *p* = 0.026, with Holm-Bonferroni correction, k = 2) and weaker Alpha oscillations (t(19) = - 2.683, *p* = 0.015, with Holm-Bonferroni correction, k = 2) for PE-open condition compared with PE-closed condition. As shown by Figure 1d, the contrast sensitivity of UPE with PE open was also higher than that with PE closed (t(17) = 2.667, *p* = 0.016).

We calculated the difference between the eye-open and eye-closed conditions as: (closed-open)/open, for all five measurements: contrast sensitivity, SSVEP amplitude, peak alpha amplitude in the SSVEP sessions (α_SSVEP_), peak alpha amplitude in the resting state when both eyes open/closed (α_Bi_) and PE open/closed (α_mono_). As shown in Figure 2, a positive correlation of the effect of eye-open v.s. eye-close was found between contrast sensitivity and SSVEP amplitude (r = 0.665, *p* = 0.013), and between peak alpha amplitudes in the SSVEP sessions and resting state (α_SSVEP_ vs. α_Bi_: r = 0.558, *p* = 0.011; α_SSVEP_ vs. α_mono_: r = 0.643, *p* = 0.002; α_Bi_ vs. α_mono_: r = 0.468, *p* = 0.037). No significant correlation was found between other pairs of measurements (all *p* > 0.05).

**Figure 2.**
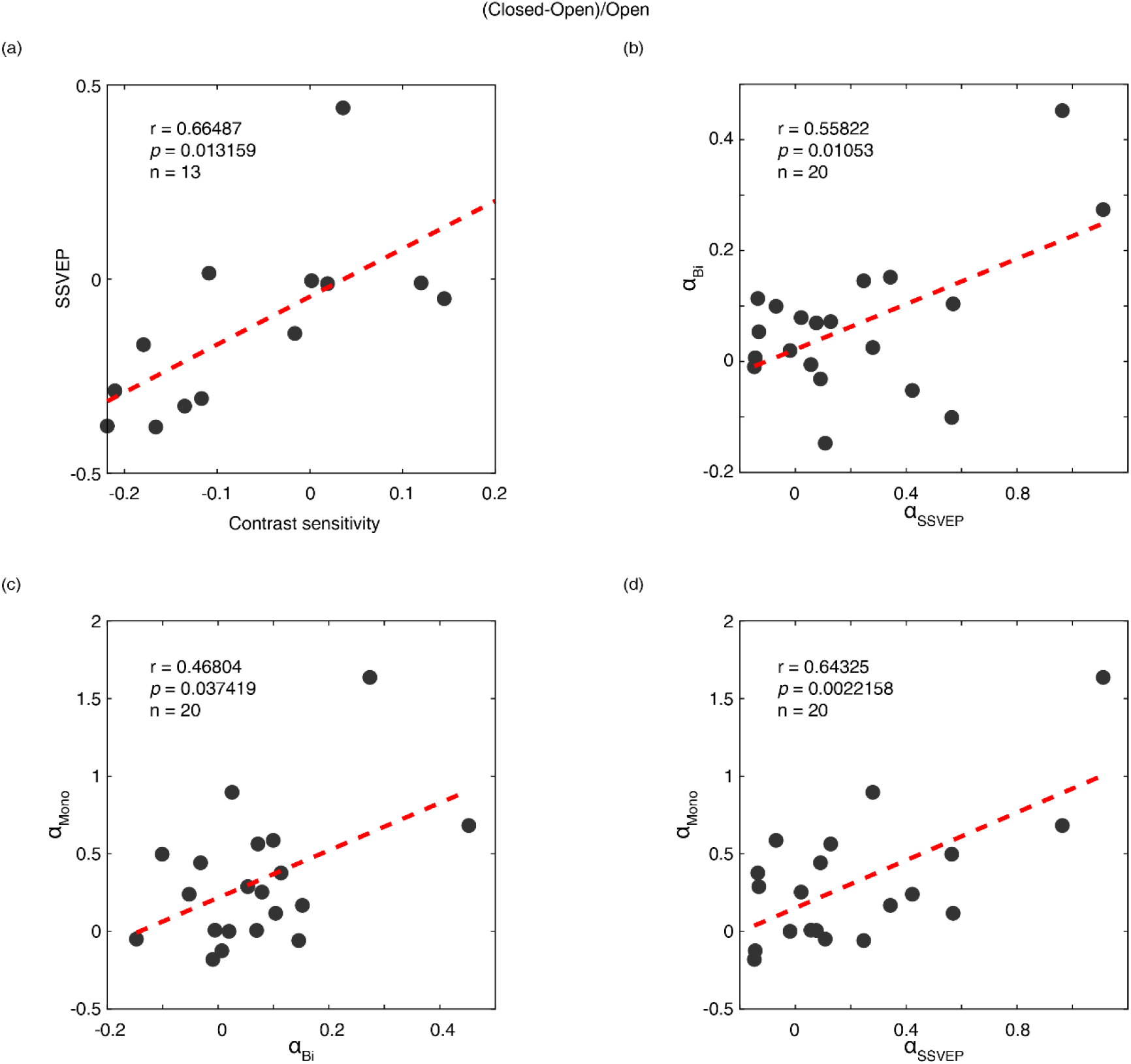
Correlations of the effects of eye-open v.s. eye-close between different measurements. The difference of the eye-open and eye-closed conditions was calculated as: (closed-open)/open. Each dot represents one observer.

### 2. The aftereffects of 2.5-hour monocular patching on contrast sensitivity

So far, we have shown that the alpha power is stronger if the eye *behind* a monocular patch is closed than when the eye *behind* the patch is open, which occurred in both the resting state and the SSVEP state. The effects of the PE closed condition were to inhibit the SSVEP power and the sensitivity of the UPE during patching.

One interesting question is whether such inhibition would influence the contrast gain changes of each eye after a short-term monocular deprivation (i.e., the aftereffects of 2.5-hour monocular patching)? In Figure 3, we plot the average change of monocular contrast sensitivity after 2.5-hour of monocular patching where the PE remains open (section M1&M3) and where the PE is kept closed (section M2&M4) as open red triangle symbols and filled black triangle symbols (UPE: dashed line with inverted triangle; PE: solid line with regular triangle), respectively. The contrast sensitivity of UPE becomes smaller for both of the two monocular deprivation conditions (for eye-closed patching, F(2,22) = 12.508, *p* < 0.001, Partial η^2^ = 0.532; for eye-open patching, F(2,22) = 19.698, *p <* 0.001, Partial η^2^ = 0.642; one-way repeated-measures within-subjects ANOVA), while the contrast sensitivity of PE becomes larger or does not change (for eye-closed patching, F(2,22) = 0.297, *p* = 0.746, Partial η^2^ = 0.026; for eye-open patching, F(2,22) = 5.212, *p* = 0.014, Partial η^2^ = 0.321; one-way repeated-measures within-subjects ANOVA).

**Figure 3.**
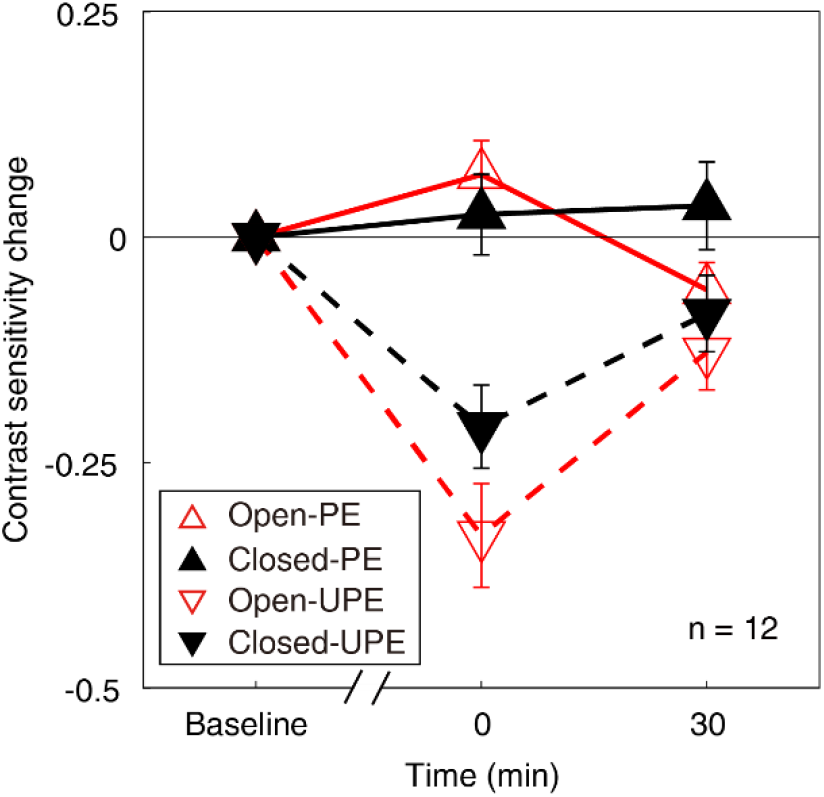
The change in monocular contrast sensitivity as a result of the monocular deprivation. The change in monocular contrast sensitivity as a result of the monocular deprivation was compared when the eye behind the patch remained open (the eye-open condition: red lines and open triangles) and when the eye behind the patch was kept closed (the eye-closed condition: black lines and filled triangles) for patched eye (solid line with regular triangle) and unpatched eye (dashed line with inverted triangle). Error bars indicate S.E.M.

We further calculated the contrast sensitivity ratio by dividing the contrast sensitivity of PE into that of UPE for two patching conditions. The change of contrast sensitivity ratio after deprivation was shown in Figure 4a. If the UPE becomes weaker or PE becomes stronger, the contrast sensitivity ratio becomes more negative, otherwise, the ratio becomes more positive. We conducted a two-way repeated-measures ANOVA, with the patching conditions (two levels) and time points of measurements after deprivation (two levels) selected as within-subject factors. The results showed that there was significant difference between two time points (F(1,11) = 20.245, *p* < 0.001, Partial η^2^ = 0.648), no difference between two patching condition(F(1,11) = 0.811, *p = 0*.*387*, Partial η^2^ = 0.069), and significant interaction of time point and condition (F(1,11) = 9.271, *p = 0*.*011*, Partial η^2^ = 0.457). Post-hoc Bonferroni test showed that the contrast sensitivity ratio changes between eye-open and eye-closed patching at 0’ was significantly different (*p = 0*.*023*, Figure 4b).

**Figure 4.**
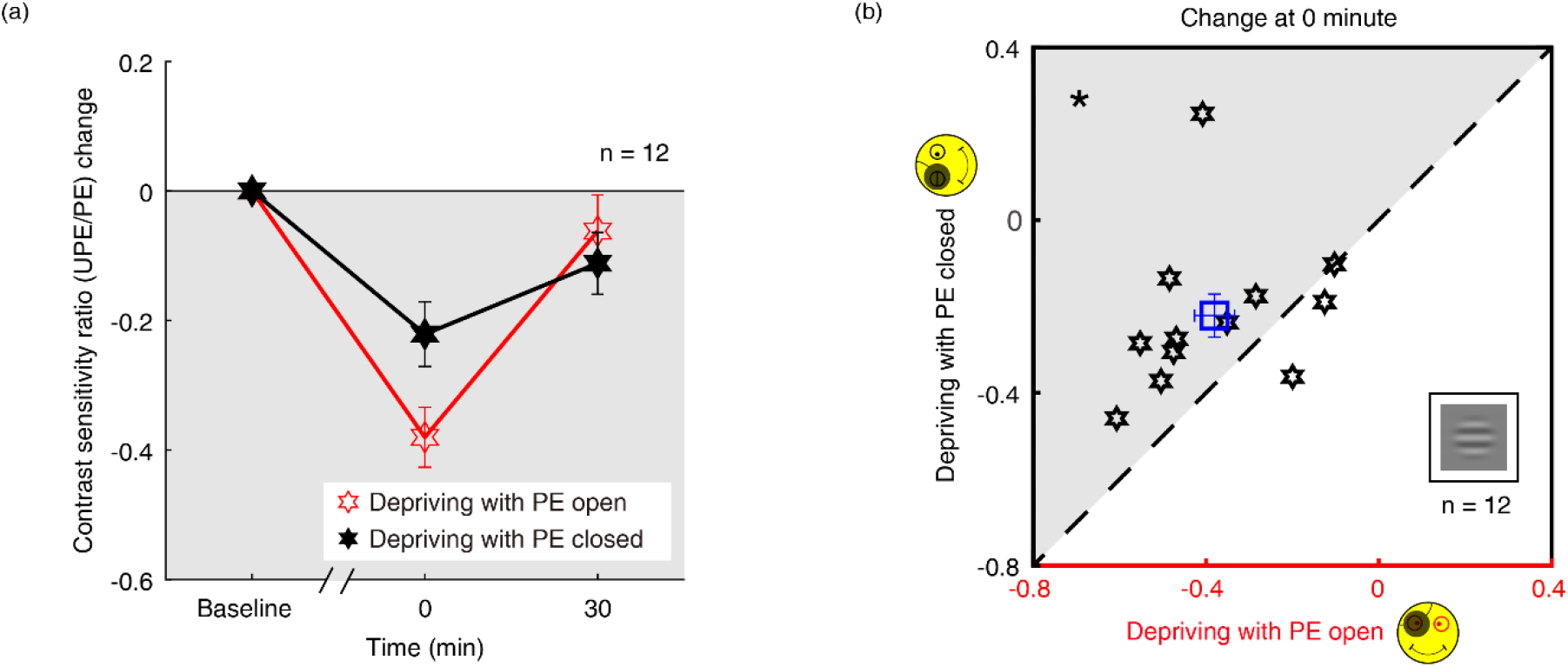
The change in contrast sensitivity ratio as a result of the monocular deprivation. (a) The change in contrast sensitivity ratio as a result of the monocular deprivation was compared when the eye behind the patch remained open (the eye-open condition: red lines and open hexagons) and when the eye behind the patch was kept closed (the eye-closed condition: black lines and filled hexagons). Error bars indicate S.E.M. (b) The average change of the post-measure session at 0’ was compared for each subject for the two patching conditions. The open square symbol represents the averaged results. The dash line is the equality line. The grey area indicates where the eye-open patching produced more patching effect than the eye-closed patching. Error bars represent standard errors across the twelve subjects.

### 3. The aftereffects of 2.5-hour monocular patching on binocular combination

Since the results show that the contrast gain changes from monocular deprivation in PE closed condition is smaller than in the PE open condition, we wondered whether such an effect could also modulate the changes of sensory eye dominance after short-term monocular deprivation? We directly tested this by monitoring changes of sensory eye dominance as a result of monocular deprivation using a binocular phase combination task. The average change of perceived phase after patching where the patched (dominant) eye remains open (section B1) and where the patched (dominant) eye is kept closed (section B2) is plotted in Figure 5a as open red square symbols and filled black square symbols, respectively. Clearly, the perceived phase changes in a more minus direction for both of the two monocular deprivation conditions. One-way repeated-measures within-subjects analysis of variance (ANOVA) showed that the binocular perceived phase significantly varied from baseline to post-measure sessions: for eye-closed patching, F(3.022,39.509) = 7.126, *p* < 0.001, Partial η^2^ = 0.354; for eye-open patching, F(5,65) = 11.420, *p* < 0.001, Partial η^2^ = 0.468. These results indicate that the PE, which was the dominant eye, was significantly strengthened after both the eye-open patching (section B1) and the eye-closed patching (section B2).

**Figure 5.**
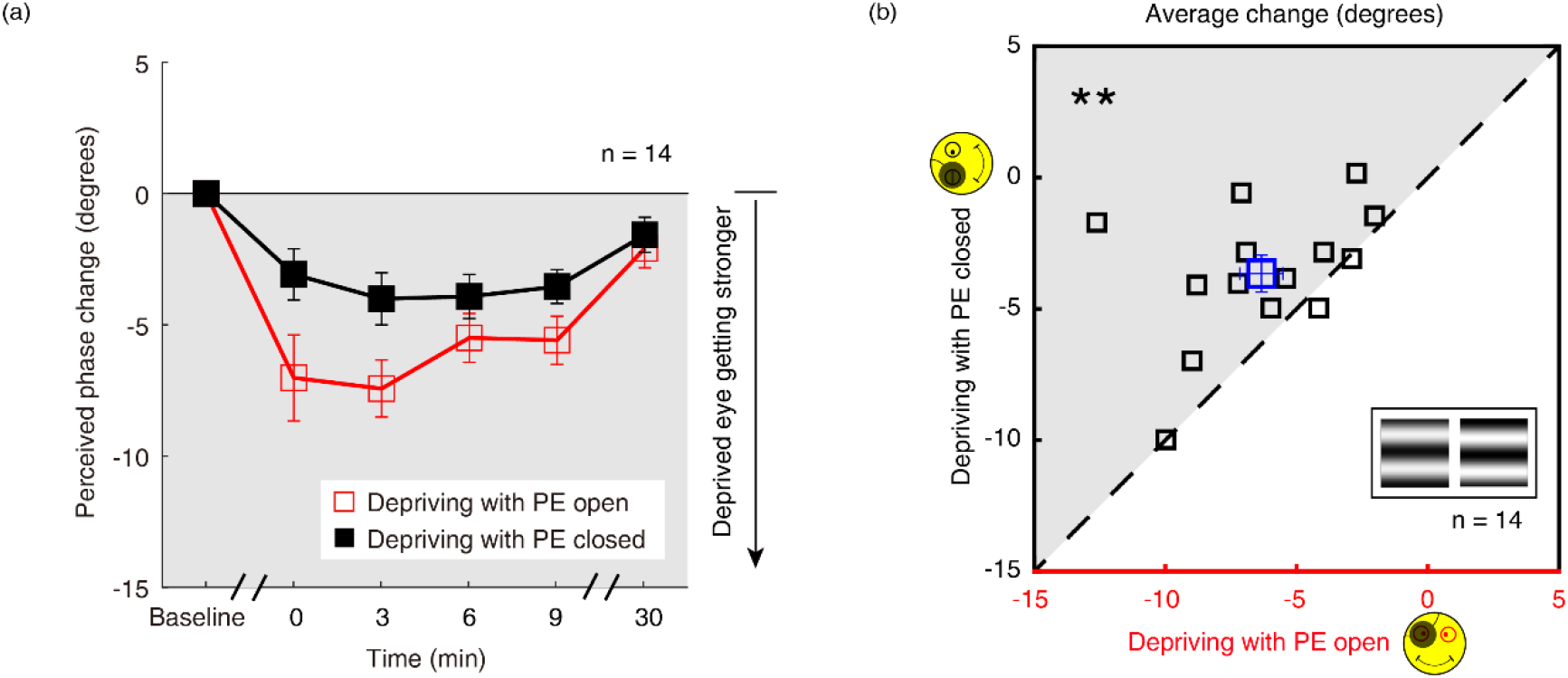
The change in ocular dominance as a result of the monocular deprivation in binocular combination. (a) The change in ocular dominance as a result of the monocular deprivation using phase combination task was compared for monocular patching where the eye behind the patch remained open (the eye-open condition-red lines and open squares) and when the eye behind the patch was closed (the eye-closed condition-black lines and filled squares). Error bars indicate standard errors across the fourteen subjects. (b) The average change (degrees) of 4 post-measure sessions (0’, 3’, 6’, 9’) using the binocular phase combination task was compared for each subject for the two patching conditions. The open square symbol represents the averaged results. The dash line is the equality line. The grey area indicates where the eye-open patching produced more cumulated shift of ocular dominance than that of the eye-closed patching. Error bars represent standard errors across the fourteen subjects.

One interesting result, which is also obvious in Figure 5a, is that the ocular dominance changes as a result of monocular deprivation are stronger in the eye-open condition than that in the eye-closed condition. A two-way repeated-measures within-subjects ANOVA also showed that the magnitude of the change of the perceived phase was significantly different between these two patching conditions (i.e., eye-open versus eye-closed): F(1,13) = 10.265, *p* = 0.007, Partial η^2^ = 0.441; the interaction between patching condition (i.e., eye-open versus eye-closed) and the post-measure sessions (i.e., from 0’ to 30’) was not significant: F(4,52) =1.553, *p* = 0.201, Partial η^2^ = 0.107, indicating the different patching impacts between eye-open and eye-closed patching was consistent during within 30 minutes after the removal of the patch.

To further show the difference between these two patching conditions, we plotted individual averages of the perceived phase change of 4 post-measure sessions (0’, 3’, 6’, 9’) with eye closed condition (section B2) as a function of that with eye-open condition (section B1) in Figure 5b. All subjects’ data, except two, located above the equality line, indicating stronger patching effect in the eye-open condition than in the eye-closed condition. A 2-tailed paired samples t-test also showed that there was a significant difference between these two conditions: t(13) = -3.276, *p* = 0.006.

### 4. The aftereffects of 2.5-hour monocular patching on binocular rivalry

Similar patching induced ocular dominance shifts were found using the binocular rivalry task (section B3&B4, Figure 6a, red and black circles) showing a distinct increase in the dominance of the PE after either eye-open patching (section B3; Figure 6a, red circles) or eye-closed patching (section B4; Figure 6a, black circles). One-way repeated-measures within-subjects ANOVA also showed that the eye dominance ratio significantly varied from baseline to post-measure sessions: for eye-closed patching, F(5,65) = 17.047, *p* < 0.001, Partial η^2^ = 0.567; for eye-open patching, F(5,65) = 34.987, *p* < 0.001, Partial η^2^ = 0.729. The magnitude of the eye dominance ratio shift was also significantly different between the two patching conditions: F(1,13) = 5.256, *p* = 0.039, Partial η^2^ = 0.288, without significant interaction between patching condition (i.e., eye-open versus eye-closed) and the post-measure sessions (i.e., from 0’ to 30’): F(4,52) =2.009, *p* = 0.107, Partial η^2^ = 0.134 (two-way repeated-measures ANOVA). In Figure 6b, the average of the perceived phase change of 4 post-measure sessions (0’, 3’, 6’, 9’) was compared for each subject for the eye-open and eye-closed patching conditions. All subjects, except three, showing stronger areal changes in the eye-open condition than in the eye-closed condition. A Wilcoxon signed-rank test also showed that there was a significant difference between two conditions: Z = -2.103, *p* = 0.035.

**Figure 6.**
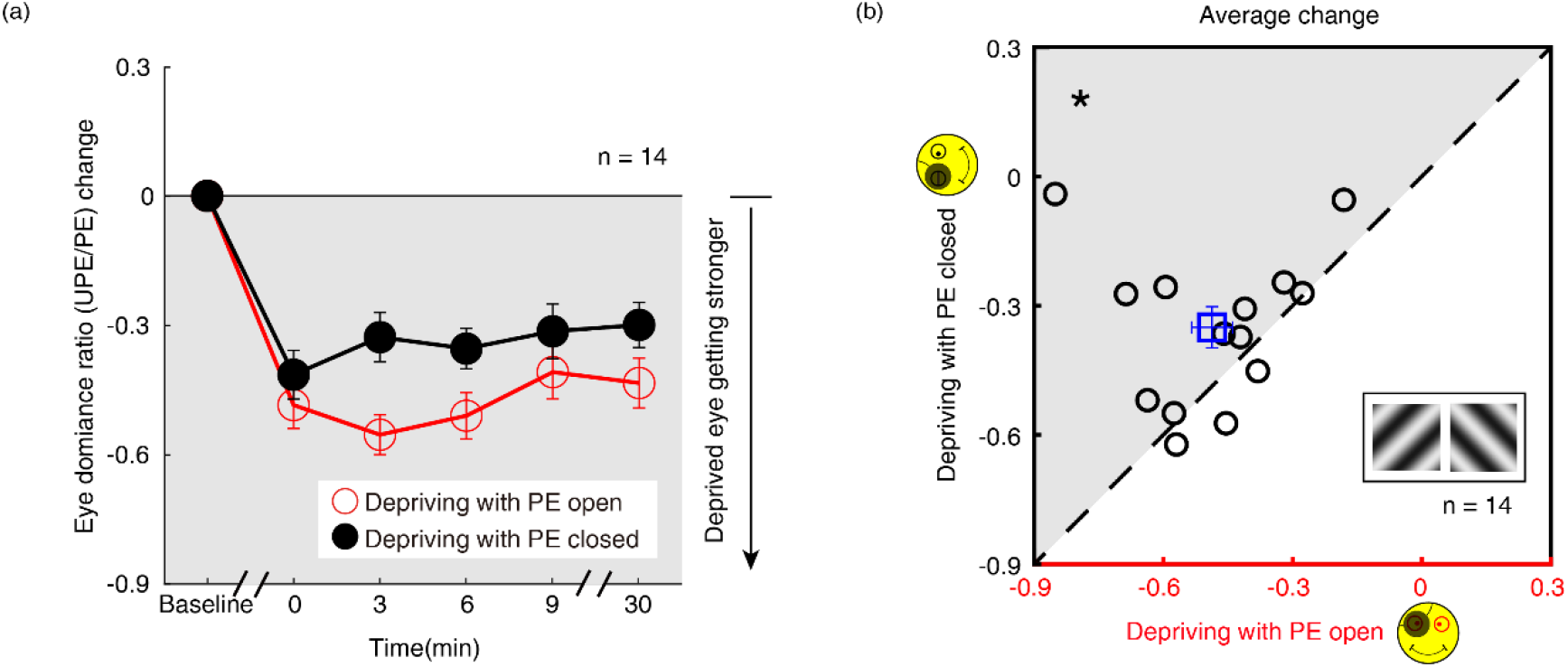
The change in ocular dominance as a result of the monocular deprivation in binocular rivalry. (a) The change in ocular dominance as a result of the monocular deprivation using binocular rivalry task was compared when the eye behind the patch remained open (the eye-open condition: red lines and open circles) and when the eye behind the patch was kept closed (the eye-closed condition: black lines and filled circles). Error bars indicate S.E.M. (b) The average change of 4 post-measure sessions (0’, 3’, 6’, 9’) using the binocular rivalry task was compared for each subject for the two patching conditions. The open square symbol represents the averaged results. The dash line is the equality line. The grey area indicates where the eye-open patching produced more cumulated shift of ocular dominance than the eye-closed patching. Error bars represent standard errors across the fourteen subjects.

## Discussion

In the absence of any visual stimulation, it is well accepted that the eyes open (REO) and eyes closed (REC) state, be it in the light or in the dark, can affect the resting state as reflected by changes in EEG spectral power and coherence in the Δ, θ, α1, α2, β1, β2 and γ frequency bands. Under complete darkness conditions, such changes cannot be related to exogenous visual stimulation, suggesting that the differences may be caused by the switching of involuntary preliminary attention from internally directed attention specific for the REC state to externally directed attention specific for the REO state (Boytsova and Danko, 2010). To our knowledge, we are the first to study the difference in brain state on visual plasticity in human adults. This was achieved by a monocular deprivation paradigm, with the patched eye (PE) that was being deprived of all visual input, being either open or closed. We show evidence that the patched eye open and closed states induce differences in alpha amplitude, which is quite similar with the differences in internal state when the *both eyes* are open versus when both eyes are closed.

The elevation of alpha oscillations when closing the PE, both for the resting state and SSVEP state, results in a reduction of contrast sensitivity of the unpatched eye (UPE) during deprivation of the other eye. This between-eye internal state effect suggests that these modulations of sensitivity by internal state manipulations are occurring at a binocular site. We also show that there is an endogenous modulation of the after-effect from short term monocular deprivation; the change in ocular dominance that results from such deprivation can be enhanced when the patched eye is kept open under the patch during the deprivation period. We show that this is true for binocular combination and also binocular rivalry even though the underlying mechanisms for these two task are thought to be very different; the former involves interocular gain control in early visual cortex (Huang, Zhou et al., 2010) and the latter involves interocular competition and top-down influences from high-level visual areas (Tong, Meng et al., 2006).

The unique aspect of the present study is that we modulate the brain’s internal state with one eye open and determine its impact on visual processing for both monocular contrast detection and binocular combination. By the manipulation of the internal states with one eye open or closed and assessing the visual sensitivity to the other eye, we can combine internal state manipulations with visual stimulation. We demonstrate, for the first time, that both visual sensitivity and its neuroplastic regulation can be modulated by endogenous factors. A finding that could have therapeutic implications. We have shown that by keeping the eye under the patch open, the resultant ocular dominance change is enhanced. If monocular patching were to be implemented as a means to rebalance the visual system of amblyopic patients (by patching the amblyopic eye in order to strengthen its contribution to binocular vision (Lunghi, Sframeli et al., 2019, Zhou, He et al., 2019)), then it would be more effective to ensure that the eye remains open under the patch. On the other hand, the more traditional therapy for amblyopia is to patch the fellow good eye to force the amblyopic eye to improve. No one has ever considered whether the eye under the patch should be open or closed. The efficacy of this approach might be improved if the child was instructed to keep their fellow eye open under the patch. This may require a redesigned patch with enough eye clearance to ensure this is possible.

The differences that we found in brain states could be explained by the contrasting dynamics of GABA, which has implications in interpreting MRS measurements (Kurcyus, Annac et al., 2018). In complete darkness, GABA concentration decreased after opening the eyes. GABA has been shown to be correlated with the sensory eye dominance plasticity, where resting GABA concentration decreases after deprivation and that the decrease in GABA correlates with the individual neuroplastic changes (Lunghi, Emir, 2015). Compared with deprivation with PE closed, deprivation with PE open would be expected to reduce the GABA levels and thus induced more change in sensory eye dominance as a result of short-term deprivation. On the other hand, previous studies show that occipital alpha inhibition originates from the pulvinar of the visual thalamus (Liu, de Zwart et al., 2012), our findings may suggest the role of pulvino-cortical inhibition in regulating ocular dominance plasticity in the early visual cortex.

Our results suggest that having the eye open under the patch, even though this does not change the exogenous stimulation because the eye is occluded, it will result in an enhanced neuroplastic effect for ocular dominance due solely to endogenous factors associated with a reduction of alpha inhibition and GABA concentration.

## Materials and Methods

### Participants

In total, fifty-one normal adults (age: 24.06 ± 2.55 years old; twenty-two males), with normal or corrected to normal vision (20/20 or better), participated in this study after having given written informed consent. All subjects were naive to the purpose of this study. In experiment 1, twenty subjects (age: 24.35 ± 2.67 years old; twelve males) participated in the EEG test, eighteen subjects (age: 24.83 ± 2.52 years old; ten males) participated in the behavioral test, and thirteen subjects (age: 24.69 ± 2.26 years old; seven males) participated in both EEG and behavioral tests. In experiment 2, twelve subjects (age: 24.08 ± 1.04 years old; three males) participated in the short-term patching study with monocular testing. In experiment 3, fourteen subjects (age: 23.21 ± 2.83 years old; four males) participated in the short-term patching study with binocular testing. Observers wore their habitual optical correction if required. The sample sizes (n ≥ 12 in all experiments) provide at least 80% power to detect a strong deprivation effect (Cohen’s d > 1) as suggested by previous studies of monocular deprivation using similar psychophysical tasks. This study complied with the Declaration of Helsinki and was approved by the Institutional Review Boards of Wenzhou Medical University. The methods were carried out in accordance with the approved guidelines.

### Apparatus

In experiment 1, EEGs were recorded using a SynAmps amplifier system (Neuroscan) with a 64 channels cap (10-20 system). Both the EEG and behavioral test were conducted using a Windows computer. The programs were written with Matlab (Mathworks, Natick, MA, United States) and PsychToolBox 3.0 (Brainard, 1997). The stimulus was presented on a CRT monitor (NESOJXC FS210A) with GAMMA corrected, of which the resolution is 2048 × 1536 pixels, and the refresh rate is 60 Hz. In experiment 2, the monocular contrast sensitivity measurement was conducted on an iMac computer using PsyKinematix software (Beaudot, 2009), and the stimuli were presented on a GAMMA corrected Built-In Retina LCD monitor (iMac, Apple, United States) in a dark room at a viewing distance of 60 cm. The monitor had a resolution of 2048 × 1152 pixels, a refresh rate of 60 Hz, and a comparable maximal luminance as goggles.

In experiment 3, sensory eye dominance measurements were conducted using a Mac computer running personally developed programs written with Matlab (Mathworks, Natick, MA, United States) and PsychToolBox 3.0 (Brainard, 1997). All stimuli were dichoptically presented using head mount goggles (Goovis pro, NED Optics, Shenzhen, China), which had a resolution of 2560 × 1600 pixels and a refresh rate of 60 Hz in each eye. The maximal luminance of the OLED goggles was 150 cd/m^2^.

## Design and procedure

### 1. EEG and behavioral tests on the immediate effects of open v.s. close of the patched eye

EEGs were recorded in six conditions for each subject: 1) patch two eyes with both eyes open (2.5 min × 2), 2) patch two eyes with both eyes closed (2.5 min × 2), 3) patch one eye with both eyes open (5 min) and keep fixation on a gray background, 4) patch one eye with the patched eye (PE) closed (5 min) and keep fixation on a gray background, 5) patch one eye with both eyes open and present flickering sine-wave plaids to the unpatched eye (UPE) (5 min), and 6) patch one eye with the PE closed and present the stimulus to the UPE (5 min). Both eyes open and closed conditions (condition 1 and 2) were tested at both the beginning and the end of the EEG experiment, with an order of “ABBA” or “BAAB” that was balanced across subjects. The dominant eye was occluded in the monocular patching conditions (condition 3-6), and a medical pressure-sensitive adhesive tape was used to help eye closure for the PE closed conditions. In condition 3 and 4, subjects were asked to keep fixation at the center of the screen on which we presented a mean luminance background of 68 cd/m^2^ (i.e., the resting state) for 5 minutes. In condition 5 and 6, a counterphase flickering sinewave plaid (2 cycles/°, 8° in diameter, 24% contrast, 7.5 Hz) was presented to the UPE to measure visually evoked signals (i.e., steady-state visual evoked potential, SSVEP). The stimulus was presented for 10 seconds for each trial, and 12 trials were collected for each condition. Subjects rested for 10 minutes in-between the PE open and closed conditions. The order of the PE open and closed conditions was counterbalanced across subjects.

EEGs were recorded at a sampling rate of 1000 Hz from the occipital and parietal electrodes, including all P, PO, O and CB channels. Electrode impedances were kept below 10 kΩ. The electrode REF on the cap between Cz and CPz was used as reference, and the electrode AFz was used as ground. EEG data were band-pass filtered from 1 to 30 Hz. For the resting state conditions (condition 1-4), EEG recordings of the first 10 seconds were removed to reduce the influence of artifacts, the remaining data were divided into 7-second epochs (i.e., same with the SSVEP-test time); for SSVEP recordings in condition 5 and 6, EEGs of the first 2 seconds and the last 1 seconds were removed for each trial. To get the peak amplitude of Alpha-band frequency, the amplitude spectrum for each epoch of each channel (occipital and parietal electrodes) was derived through fast Fourier transform (FFT) and was then averaged across epochs and channels. Then, Savitzky-Golay filter was performed on the averaged spectrum, and the peak value between 8 and 13 Hz was taken as the peak amplitude of Alpha-band oscillation. To get the amplitude of SSVEP signals, we first averaged the EEG data across all trials and channels, and then derived the amplitude spectrum with FFT. The amplitude of the second harmonic at 15 Hz was taken as the SSVEP amplitude. To reduce the influence of large variations of SSVEP amplitude across subjects on the statistical results, the amplitude of SSVEPs of all conditions for each subject were normalized (divided) by the mean SSVEP amplitude across all conditions. The normalized amplitudes were then multiplied by the mean amplitudes across all subjects. Alpha amplitudes were normalized in the same way.

In a behavioral test, we measured contrast sensitivity for the UPE when the dominant eye was occluded. Eighteen observers, 13 of whom were the same as the EEG test, participated in the behavioral measurement. The behavioral test contains two conditions: the PE-open condition and the PE-close condition. The order of two conditions was counterbalanced across subjects. In this test, the contrast sensitivity was measured using an adjustment method. In particular, a 45° or 135° oriented sinewave grating (2 cycles/°, 8° in diameter) was presented to the UPE with an initial contrast lower than the observer’s threshold. During the measurement, subjects were instructed to adjust the contrast of the grating by key pressing until they were just able to discriminate the orientation of the grating, and this contrast level was taken as the contrast threshold. Contrast sensitivity was defined as the inverse of contrast threshold. There were 40 trials for each condition. Subjects practiced 40-60 trials before the formal test.

### 2. The aftereffect of short-term patching on monocular contrast sensitivity

Twelve new observers participated in four patching sections. As with the psychophysical binocular test, each patching section consisted of three consecutive stages: a pre-deprivation measurement of contrast sensitivity (baseline), a 2.5-hour monocular deprivation stage and a post-deprivation measurement of contrast sensitivity. For each observer, the dominant eye (assessed by the hole-in-the-card test; (Dane and Dane, 2004)) was chosen for short-term monocular deprivation. Observers were free to do any visual work, except sleeping or exercising under patching. During the monocular deprivation stage, observers’ dominant eye was patched by a black patch with the PE open in section M1&M3 and with the PE closed in section M2&M4 (a medical pressure-sensitive adhesive tape was also used to ensure observers’ PE was closed). Different deprivation conditions were conducted in a random order on different days for different observers.

An orientation discrimination task (Figure 7a) was used to measure monocular contrast sensitivity of PE (section M1&M2) and UPE (section M3&M4). In this task, the monocular contrast function was measured using a constant stimuli method. In particular, a vertical or horizontal oriented sine-wave grating (0.46 cycle/°, 4.33°×4.33°) was presented to the PE (section M1&M2) or UPE (section M3&M4). During the measurement, the unmeasured eye was patched by an opaque patch. Subjects were instructed to answer whether the orientation of the grating was vertical or horizontal by key pressing. Probabilities of correct identification were measured at six contrast levels; each level contains 75 trials, lasting for 15 minutes. We fitted the psychometric functions using Quick functions with parametric maximum likelihood estimation (Watson, 1979). The parameter alpha of the Quick function represents the threshold corresponding to 81.6% accuracy, whose mean and variances were determined by 500 times’ bootstrap simulation (Efron and Tibshirani, 1994). Contrast sensitivity was defined as the inverse of contrast threshold. Using this approach, observers’ monocular contrast sensitivity of each eye was assessed before the deprivation and at 0’ and 30’ after the completion of the 2.5-hour of monocular deprivation.

**Figure 7.**
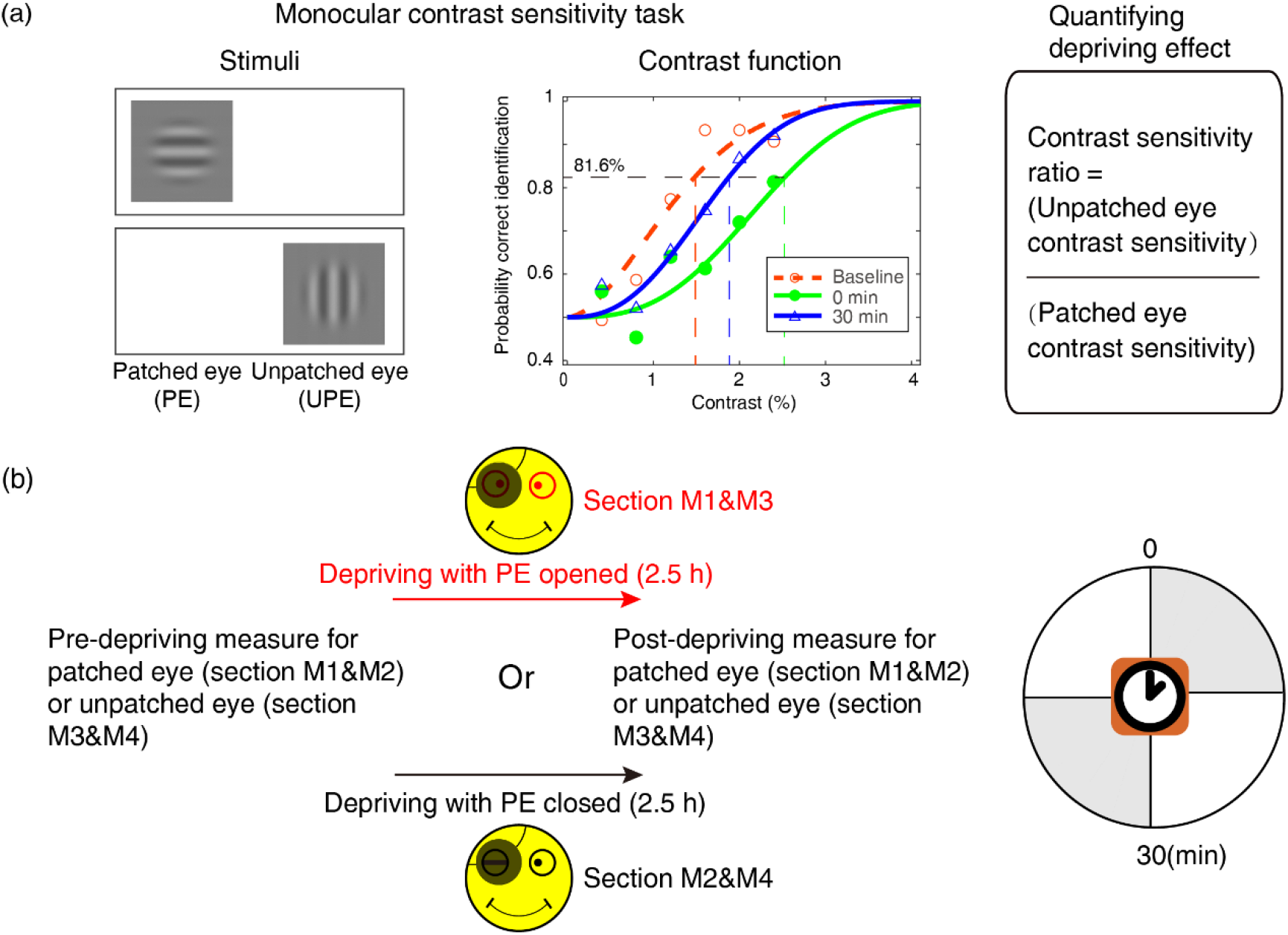
Experimental design and procedure of psychophysical monocular test. (a) The monocular contrast sensitivity task. The stimulus was a vertical or horizontal sine-wave grating, which was presented to the patched eye or unpatched eye with the unmeasured eye occluded. Observers’ contrast response functions were measured with constant stimuli method. Quick functions with maximum likelihood estimation were used to fit the contrast response function and derive the contrast thresholds. Contrast sensitivity was defined as the inverse of contrast threshold. Patching effect was quantified by the change of interocular contrast sensitivity ratio. (b) Monocular contrast sensitivity was measured before and after the 2.5-hour patching stage for patched eye or unpatched eye, started at multiple time points (0-30 min) after eye-open patching or eye-closed patching removal.

### 3. The aftereffect of short-term patching on binocular imbalance

Fourteen observers participated in another four patching sections to measure the effects of monocular patching on binocular imbalance. Each patching section also consisted of three consecutive stages: a pre-deprivation measurement of sensory eye balance (baseline), a 2.5-hour monocular deprivation stage and a post-deprivation measurement of sensory eye balance. For each observer, the dominant eye (assessed by the hole-in-the-card test; (Dane and Dane, 2004)) was chosen for short-term monocular deprivation. Observers were free to do any visual work, except sleeping or exercising under patching. During the monocular deprivation stage, observers’ dominant eye was patched by a black patch with the patched eye (PE) open in section B1&B3 and with the PE closed in section B2&B4 (a medical pressure-sensitive adhesive tape was also used to ensure observers’ PE was closed). Different deprivation conditions were conducted in a random order on different days for different observers.

In section B1&B2, a binocular phase combination task (Zhou, Clavagnier, 2013) was used to quantitatively assess the degree of sensory eye balance, in which the binocularly perceived phase was measured and used as an index of sensory eye dominance (Figure 8a). In particular, two horizontal sine-wave gratings (0.46 cycle/°, 4.33°×4.33°) with equal and opposite phase-shift of 22.5° relative to the center screen were dichoptically presented to the two eyes; the contrast of the unpatched eye (UPE) was fixed at 100% and the contrast of the PE was chosen close to an individuals’ balance point in binocular phase combination before the deprivation (at that balance point, observer’s two eyes are equally effective in binocular phase combination and the binocular perceptive phase is 0°). Two configurations were used to cancel any potential positional bias. In configuration 1, the phase-shift was +22.5° from horizontal in the UPE and -22.5° from horizontal in the PE and in configuration 2, the reverse. Each configuration was repeated eight times in one measurement session, in which 16 trials (8 repetitions × 2 configurations) were randomly interleaved. Normally, subjects could finish one measurement in three minutes after a short period of practice. Observers’ binocular perceived phase was calculated by the averaged difference between the two configurations and was assessed before deprivation and at 0’, 3’, 6’, 9’ and 30’ after the completion of the 2.5-hour of monocular deprivation. Thus, if the PE became stronger, the binocular perceived phase would be decreased, otherwise, if the PE became weaker, the binocular perceived phase would be increased.

**Figure 8.**
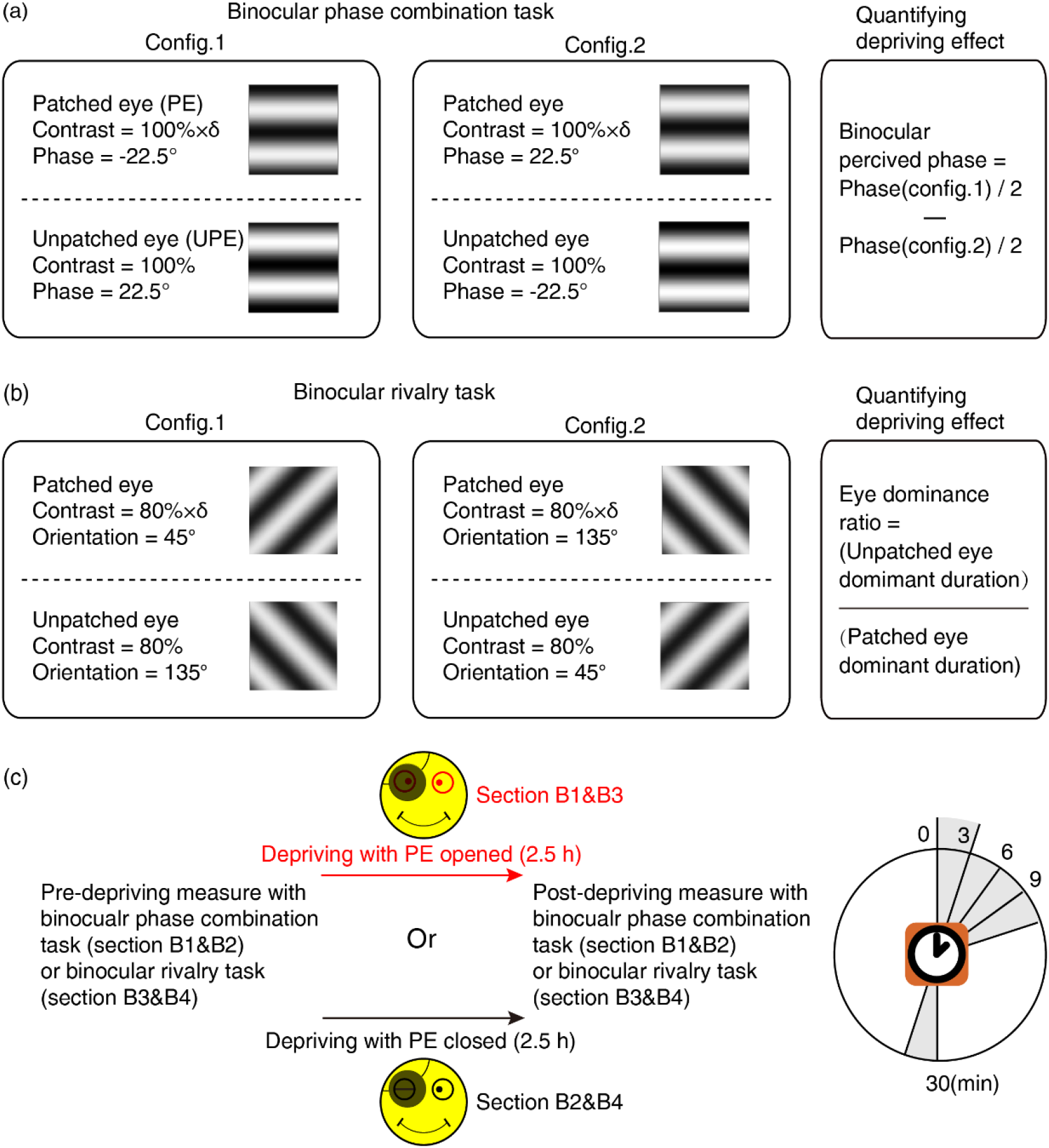
Experimental design and procedure of psychophysical binocular test. (a) The binocular phase combination task. The stimuli were two horizontal sine-wave gratings with equal and opposite phase-shift of 22.5° relative to the horizontal center of the screen, which were dichoptically presented to the two eyes. Patching effect on sensory eye dominance was quantified by the change of binocularly perceived phase. (b) The binocular rivalry task. The stimuli were two orthogonal sine-wave gratings, which were dichoptically presented to the two eyes. Patching effect on sensory eye dominance was quantified by the change of eye dominance ratio. (c) Sensory eye dominance was measured before and after the 2.5-hour patching stage by phase combination task or rivalry task, started at multiple time points (0-30 min) after eye-open patching or eye-closed patching removal.

In section B3&B4, a binocular rivalry task was used to quantitatively assess the sensory eye balance. In this task, the interocular ratio of total phase duration (UPE / PE) was measured and used as an index of sensory eye dominance (Figure 8b). In particular, two vertically orthogonally oriented sine-wave gratings (0.46 cycle/°, 4.33°×4.33°) were dichoptically presented to the two eyes; the contrast of the nondominant eye was fixed at 80%, and the contrast of the dominant eye was chosen close to an individuals’ balance point in binocular rivalry before deprivation (at that balance point, observer’s two eyes are equally effective in binocular rivalry and the ratio of total phase duration closed to 1). Each test session consists of two 90-second sub-blocks for two configurations: in configuration1, the orientation was 135° in the UPE and was 45° in the PE; in configuration 2, the orientation was 45° in the UPE and was 135° in the PE. Using this approach, observers’ eye dominance ratio was calculated by the ratio between the UPE dominant duration and the PE dominant duration, and assessed before the deprivation and at 0’, 3’, 6’, 9’ and 30’ after the completion of the 2.5-hour of monocular deprivation. Thus, if the PE became stronger, the sensory eye dominance ratio became more negative, otherwise, the ratio became more positive.

### Statistical Analysis

SPSS v.23.0 (IBM Corporation, Armonk, NY, USA) was used for statistical analyses. We compared the immediate effects on EEG and behaviors between the PE open and closed condition using a 2-tailed paired samples t-test with Holm-Bonferroni correction(Holm, 1979). We also conducted a Pearson correlation test to find the relationship between the effects on EEG and behaviors. Repeated-measures within-subjects ANOVA was applied to evaluate the effect of time after deprivation and the patching conditions (i.e., PE open and PE closed) on contrast sensitivity and binocular balance.

## Data Availability

All data generated or analyzed during this study are included in the manuscript.

## Author Contributions

J.Z., P.Z., and R.F.H. designed research; Y.C., Y.G., Z.H., Z.S. and Y.M. performed research; Y.C., Y.G. and Z.H. analyzed data; J.Z. and R.F.H. acquired funding; and J.Z., P.Z., R.F.H., Y.C., Y.G. and Z.H. wrote the paper.

## Acknowledgments

This work was supported by the National Natural Science Foundation of China Grant (NSFC 31970975), the Natural Science Foundation for Distinguished Young Scholars of Zhejiang Province, China (LR22H120001), and the Project of State Key Laboratory of Ophthalmology, Optometry and Vision Science, Wenzhou Medical University (No. J02-20210203) to JZ, the Canadian Institutes of Health Research Grants CCI-125686, NSERC grant 228103, and an ERA-NET Neuron grant (JTC2015) to RFH. The sponsor or funding organization had no role in the design or conduct of this research.

## References

Beaudot W. (2009) Psykinematix: A new psychophysical tool for investigating visual impairment due to neural dysfunctions. Journal of the Vision Society of Japan 21:19–32.

Begum M, Tso DY. (2015) Short-term monocular deprivation reveals rapid shifts in interocular balance and gain in adult macaque visual cortex. Invest Ophthalmol Vis Sci 56(7).

Binda P, Kurzawski JW, Lunghi C, Biagi L, Tosetti M, Morrone MC. (2018) Response to short-term deprivation of the human adult visual cortex measured with 7T BOLD. Elife 7. doi:10.7554/eLife.40014.

Boytsova YA, Danko SG. (2010) EEG differences between resting states with eyes open and closed in darkness. Hum Physiol 36(3):367–9.

Brainard DH. (1997) The Psychophysics Toolbox. Spat Vis 10(4):433–6. doi:10.1163/156856897x00357

Brodoehl S, Klingner CM, Witte OW. (2015) Eye closure enhances dark night perceptions. Sci Rep 5:10515. doi:10.1038/srep10515.

Chadnova E, Reynaud A, Clavagnier S, Hess RF. (2017) Short-term monocular occlusion produces changes in ocular dominance by a reciprocal modulation of interocular inhibition. Sci Rep 7. doi:10.1038/srep41747.

Dane A, Dane S. (2004) Correlations among handedness, eyedness, monocular shifts from binocular focal point, and nonverbal intelligence in university mathematics students. Percept Mot Skills 99(2):519–24. doi:10.2466/Pms.99.6.519-524.

Efron B, Tibshirani RJ. An Introduction to the Bootstrap (Vol. 57). New York: Chapman & Hall/CRC; 1994.

Holm S. (1979) A simple sequentially rejective multiple test procedure. Scandinavian Journal of Statistics 6(2):65–70.

Huang CB, Zhou J, Zhou Y, Lu ZL. (2010) Contrast and phase combination in binocular vision. PLoS One 5(12):e15075. doi:10.1371/journal.pone.0015075.

Kurcyus K, Annac E, Hanning NM, Harris AD, Oeltzschner G, Edden R, Riedl V. (2018) Opposite Dynamics of GABA and Glutamate Levels in the Occipital Cortex during Visual Processing. J Neurosci 38(46):9967–76. doi:10.1523/JNEUROSCI.1214-18.2018.

Liu Z, de Zwart JA, Yao B, van Gelderen P, Kuo LW, Duyn JH. (2012) Finding thalamic BOLD correlates to posterior alpha EEG. Neuroimage 63(3):1060–9. doi:10.1016/j.neuroimage.2012.08.025.

Lunghi C, Berchicci M, Morrone MC, Di Russo F. (2015) Short-term monocular deprivation alters early components of visual evoked potentials. J Physiol 593(19):4361–72. doi:10.1113/Jp270950.

Lunghi C, Burr DC, Morrone C. (2011) Brief periods of monocular deprivation disrupt ocular balance in human adult visual cortex. Curr Biol 21(14):R538–R9. doi:10.1016/j.cub.2011.06.004.

Lunghi C, Daniele G, Binda P, Dardano A, Ceccarini G, Santini F, Del Prato S, Morrone MC. (2019) Altered Visual Plasticity in Morbidly Obese Subjects. iScience 22:206–13. doi:10.1016/j.isci.2019.11.027.

Lunghi C, Emir UE, Morrone MC, Bridge H. (2015) Short-Term Monocular Deprivation Alters GABA in the Adult Human Visual Cortex. Curr Biol 25(11):1496–501. doi:10.1016/j.cub.2015.04.021.

Lunghi C, Galli-Resta L, Binda P, Cicchini GM, Placidi G, Falsini B, Morrone MC. (2019) Visual Cortical Plasticity in Retinitis Pigmentosa. Invest Ophthalmol Vis Sci 60(7):2753–63. doi:10.1167/iovs.18-25750.

Lunghi C, Sale A. (2015) A cycling lane for brain rewiring. Curr Biol 25(23):R1122–R3. doi:10.1016/j.cub.2015.10.026.

Lunghi C, Sframeli AT, Lepri A, Lepri M, Lisi D, Sale A, Morrone MC. (2019) A new counterintuitive training for adult amblyopia. Ann Clin Transl Neur 6(2):274–84. doi:10.1002/acn3.698.

Marx E, Deutschlander A, Stephan T, Dieterich M, Wiesmann M, Brandt T. (2004) Eyes open and eyes closed as rest conditions: impact on brain activation patterns. Neuroimage 21(4):1818–24. doi:10.1016/j.neuroimage.2003.12.026.

Min SH, Chen Y, Jiang N, He Z, Zhou J, Hess RF. (2022) Issues Revisited: Shifts in Binocular Balance Depend on the Deprivation Duration in Normal and Amblyopic Adults. Ophthalmol Ther. doi:10.1007/s40123-022-00560-5.

Tong F, Meng M, Blake R. (2006) Neural bases of binocular rivalry. Trends Cogn Sci 10(11):502–11. doi:10.1016/j.tics.2006.09.003.

Watson AB. (1979) Probability summation over time. Vision Res 19(5):515–22. doi:10.1016/0042-6989(79)90136-6.

Zhou J, Clavagnier S, Hess RF. (2013) Short-term monocular deprivation strengthens the patched eye’s contribution to binocular combination. J Vision 13(5):1–10. doi:10.1167/13.5.12.

Zhou J, He Z, Wu Y, Chen Y, Chen X, Liang Y, Mao Y, Yao Z, Lu F, Qu J, Hess RF. (2019) Inverse Occlusion: A Binocularly Motivated Treatment for Amblyopia. Neural Plast 2019:5157628. doi:10.1155/2019/5157628.

Zhou J, Reynaud A, Hess RF. (2014) Real-time modulation of perceptual eye dominance in humans. Proc Biol Sci 281(1795). doi:10.1098/rspb.2014.1717.

Zhou J, Reynaud A, Kim YJ, Mullen KT, Hess RF. (2017) Chromatic and achromatic monocular deprivation produce separable changes of eye dominance in adults. Proc Biol Sci 284(1867). doi:10.1098/rspb.2017.1669.

Zhou JW, Baker D, Simard M, Saint-Amour D, Hess RF. (2015) Short-term monocular patching boosts the cortical response to the patched eye. Invest Ophthalmol Vis Sci 56(7).

